# FEATMAP: Targeted Correction of Acquisition Signatures Harmonizes Medical Foundation Model Embeddings and Enables Robust Task Generalization

**DOI:** 10.64898/2026.07.02.736184

**Authors:** Leonhard Donle, Michael Phillips, Farieda Gaber, Siddhi Ramesh, Matteo Sacco, Sampsa Hautaniemi, Anni Virtanen, Keno Bressem, Lisa Adams, Kelsey Goon, Elena Nevins, Ryan A. Robinett, Sara Kochanny, Sasha Hassan, James Dolezal, Alexander T. Pearson, Ernst Lengyel

**Author notes:** Correspondence: Ernst Lengyel; Alexander T. Pearson. These authors contributed equally: Leonhard Donle and Michael Phillips.

## Abstract

Medical foundation models compress biomedical data into embeddings that support diverse downstream clinical tasks. However, successful model deployment is hampered by performance degradation on external data. It is recognized that embeddings capture acquisition signatures, such as hardware and technical differences, in addition to biology. Effective harmonization must remove the acquisition signature while preserving biological signals, a trade-off that current methods fail to balance adequately. Input-level normalization fails to eliminate acquisition signatures from embeddings, whereas embedding-level methods adjust features in an untargeted manner. We present FEATMAP, a harmonization approach that models acquisition signatures as geometric distortions between manifolds of similarly arranged embeddings. Using paired data that isolate the effect of acquisition signatures, FEATMAP fits a single global affine transformation per foundation model to correct acquisition signatures directly in the embedding space. This targeted, reusable correction aims to preserve biological and demographic variation while harmonizing across acquisition signatures. Across scanner and foundation-model harmonization in digital pathology and field-strength harmonization in brain MRI, FEATMAP improves cross-condition embedding similarity, reduces performance gaps without retraining, and suggests potential for the alignment of disparate embedding spaces.

## 1 Introduction

The transformative success of medical foundation models comes from their ability to learn biologically meaningful representations from large, unlabeled datasets across many medical modalities, including histopathology [1] and radiology [2, 3]. These foundation models are widely used as feature extractors, compressing complex biomedical inputs into high-dimensional numerical vectors termed *embeddings*. These embeddings support downstream fine-tuning across a broad range of clinical tasks, help reduce bias in fine-tuned downstream models trained on modest-sized datasets, and decrease reliance on dense expert annotation.

However, reliable deployment across institutions and cohorts remains difficult because foundation model embeddings often capture acquisition and processing signatures unrelated to biology [4]. Metadata leakage occurs when variation of these domains leaks into the embeddings and produces separable feature distributions across datasets even when the underlying biology is unchanged [5, 6]. We refer to these non-biological, domain-specific signals as acquisition signatures, as they are systematic patterns induced by acquisition conditions such as scanner hardware and staining protocol in pathology, magnetic field strength in magnetic resonance imaging, and digitization and learning architecture biases, among others [7]. In computational pathology, individual sites that submit tissue to The Cancer Genome Atlas (TCGA) can be identified based on site characteristics in the images despite their lack of biological relevance [8], and the same tissue digitized on different scanners yields systematically different embeddings that hinder cross-cohort comparison. In radiology, MRI scanners operating at different field strengths introduce acquisition-driven intensity and contrast differences that similarly propagate into foundation model embeddings, degrading cross-site generalization. This leakage can artificially inflate apparent performance when site-specific signals correlate with patient demographics and outcomes [8] and can substantially degrade performance on external validation datasets where a domain shift is present [9, 10].

Prior work has attempted to mitigate domain shifts at the input level, before feature extraction, through modality-specific normalization [7, 11]. These include stain normalization in pathology, such as Reinhard [12] and Macenko [13], or intensity normalization in MRI, such as histogram equalization [14]. However, despite aligning visible appearance these methods do not reliably eliminate separable domain signatures [8], and reported improvements in external generalization remain limited or inconsistent [9, 15]. Existing embedding-level harmonization methods address the problem directly where downstream prediction models operate, but also carry fundamental limitations. First, many methods treat the embedding space as a collection of independent features, applying coordinate-wise corrections that adjust each dimension separately under simple distributional models [16, 17]. This assumption is poorly suited to foundation model embeddings because, in contrast to omics-style data, where individual axes can often correspond to interpretable biological quantities, foundation model embeddings encode information jointly across embedding dimensions rather than in individual coordinates. Consistent with this, evidence suggests that high-level structure in embedding spaces is not confined to simple linear axes [18] and that embeddings can exhibit nonlinear and radial organization [19, 20]. Second, existing methods are non-targeted: they adjust features irrespective of the source of variation and correct any structural differences simultaneously, thereby risking the removal of relevant biological variation alongside acquisition signatures. Finally, many approaches are fitted jointly with a downstream prediction objective and therefore require retraining for each new task or cohort, preventing deployment as reusable harmonization modules [21, 22]. To our knowledge, only one existing approach addresses all of these limitations, but it relies on aligned language information in the foundation model to guide acquisition signature suppression [23]. This requirement limits its applicability to vision-language models, whereas most medical foundation models lack such aligned language representations.

To address these limitations, we introduce Feature Embedding Affine Transformations for foundation Model Agnostic Projections (FEATMAP), a framework that learns a targeted, reusable transformation between acquisition domains directly in the embedding space to harmonize acquisition signatures (Fig. 1). FEATMAP was trained on paired data in which biological content is held constant while only the acquisition signature is systematically varied, so that the learned transformation reflects the effect of the different acquisition signatures alone, disentangled from biological variability.

**Fig. 1.**
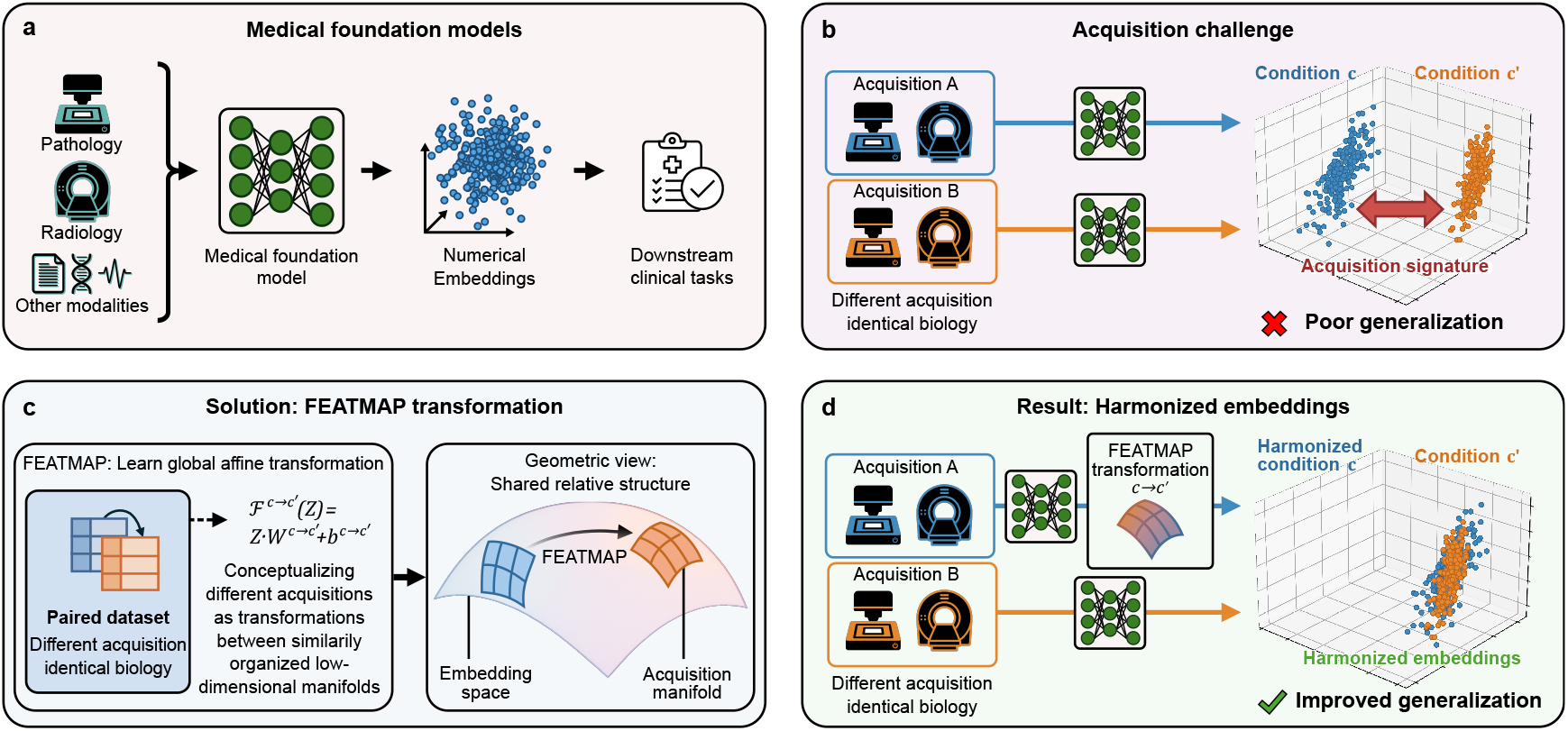
| Overview of the FEATMAP framework. **a**, Medical foundation models serve as universal feature extractors across biomedical modalities including pathology, radiology, and others, by compressing biomedical inputs into high-dimensional numerical embeddings for downstream clinical tasks. **b**, The acquisition challenge arises when the same biological content is acquired under different acquisition conditions (e.g., different pathological scanners or MRI machines) and the resulting embeddings separate by acquisition signature rather than biology, producing an acquisition signature that impairs cross-dataset generalization. **c**, FEATMAP addresses this challenge by modeling domain shift as a geometric distortion between similarly organized lower-dimensional manifolds in the embedding space. Using paired data in which biological content is held constant while the acquisition signature varies, FEATMAP learns a single global affine transformation that maps embeddings from one acquisition signature to another. **d**, The resulting FEATMAP harmonized embeddings reduce acquisition-driven separation while preserving the biological signal, enabling robust generalization for medical downstream tasks across different acquisition signatures.

Conceptually, FEATMAP treats shifts in the acquisition domain as a geometric distortion between lower-dimensional manifolds of the full embedding space. In this view, the embeddings produced under each acquisition signature occupy their own manifold with a shared local structure within the embedding space. This allows movement between signatures by reshaping one manifold onto the other. This view is consistent with the Manifold Hypothesis [24], the Union of Manifolds Hypothesis [25] and Dvoretzky’s theorem [26]. Together, these propose that high-dimensional embedding spaces contain collections of lower-dimensional surfaces that are locally linearly disentangled [20, 27] and behave approximately Euclidean [22]. It is further supported by the Platonic Representation Hypothesis [28], which suggests that models trained to represent real-world data preserve underlying causal structure, making it likely that acquisition signatures occupy two similarly shaped manifolds within the embedding space. These geometric assumptions are not imposed explicitly by design, but instead emerge implicitly from the training objectives of many foundation models, as contrastive learning and self-distillation rely on attractor–repulsor dynamics that shape the embedding space by enforcing geometric constraints aligned with semantic relationships [29].

Under these premises, two acquisition signatures correspond to two manifolds with a shared geometric structure. A complete reorganization of the embedding space through a nonlinear mapping between them (as commonly done in methods applying optimal transport [30]) is therefore unnecessary and introduces far more degrees of freedom than the problem requires. The most principled and efficient correction is instead a geometric transformation, specifically an affine transformation, which captures translation, rotation, scaling, and shearing. FEATMAP fits a single global affine transformation per foundation model, enabling embeddings from one acquisition condition to be mapped into another. Under our assumptions, FEATMAP aims to preserve all remaining variation, including biological and demographic structure, that does not reflect the targeted acquisition signature.

The same geometric reasoning also applies across foundation model embedding spaces. Since the Platonic Representation Hypothesis [28] proposes that embeddings converge toward a shared statistical description of reality, foundation models trained on overlapping histopathology data may encode a common embedding structure reflecting shared biological morphology, even though the resulting embedding spaces are not directly comparable. We hypothesize that FEATMAP can utilize this shared geometric structure between foundation models to learn affine mappings between foundation model embedding spaces using the same paired-data strategy applied to acquisition harmonization, instead of a complete reorganization as typically performed [31].

Herein, we show that FEATMAP generalizes across three distinct settings for foundation model acquisition harmonization: scanner harmonization in digital pathology, foundation model harmonization in digital pathology, and field-strength harmonization in brain MRI. In each setting, we trained FEATMAP on a paired discovery dataset and evaluated generalization on an independent, external, public validation dataset. Across all settings, FEATMAP substantially increases cross-signature embedding similarity and reduces downstream performance gaps between acquisition signatures, without requiring retraining for each new task or dataset. FEATMAP thereby provides a scalable, modular, and targeted approach to correct specific acquisition signatures in medical foundation model embeddings. This strengthens comparability between acquisition protocols, allowing the creation of larger cross-institutional datasets, and enables reliable deployment in real-world clinical settings.

## 2 Results

### 2.1 FEATMAP harmonizes across the digital pathology scanner domain

The same tissue digitized on different scanners produces systematically different foundation model embeddings, even though the underlying biology is unchanged. To address this, we trained FEATMAP on the discovery dataset from the University of Chicago (UChicago) which comprises approximately 8.4 million pixel-perfect aligned tile embeddings across four whole-slide-image scanners (AT2, VS200, GT450 and O40) and five foundation models (UNI [35], H-Optimus-0 [36], Conch [37], Prov-GigaPath [38] and Virchow2 [39]). The learned mapping was then evaluated on the validation dataset PLISM [40, 41], which contains approximately 10.6 thousand matched tiles across two scanners, 46 tissue types and three staining protocols. To provide an unbiased comparison, FEATMAP was benchmarked against five state-of-the-art harmonization methods spanning stain normalization (Macenko [12], Reinhard [13]), stain transfer (StainGAN [32], CAGAN [33]) and embedding-level correction (ComBat [34]), as well as an uncorrected baseline (Fig. 2a).

**Fig. 2.**
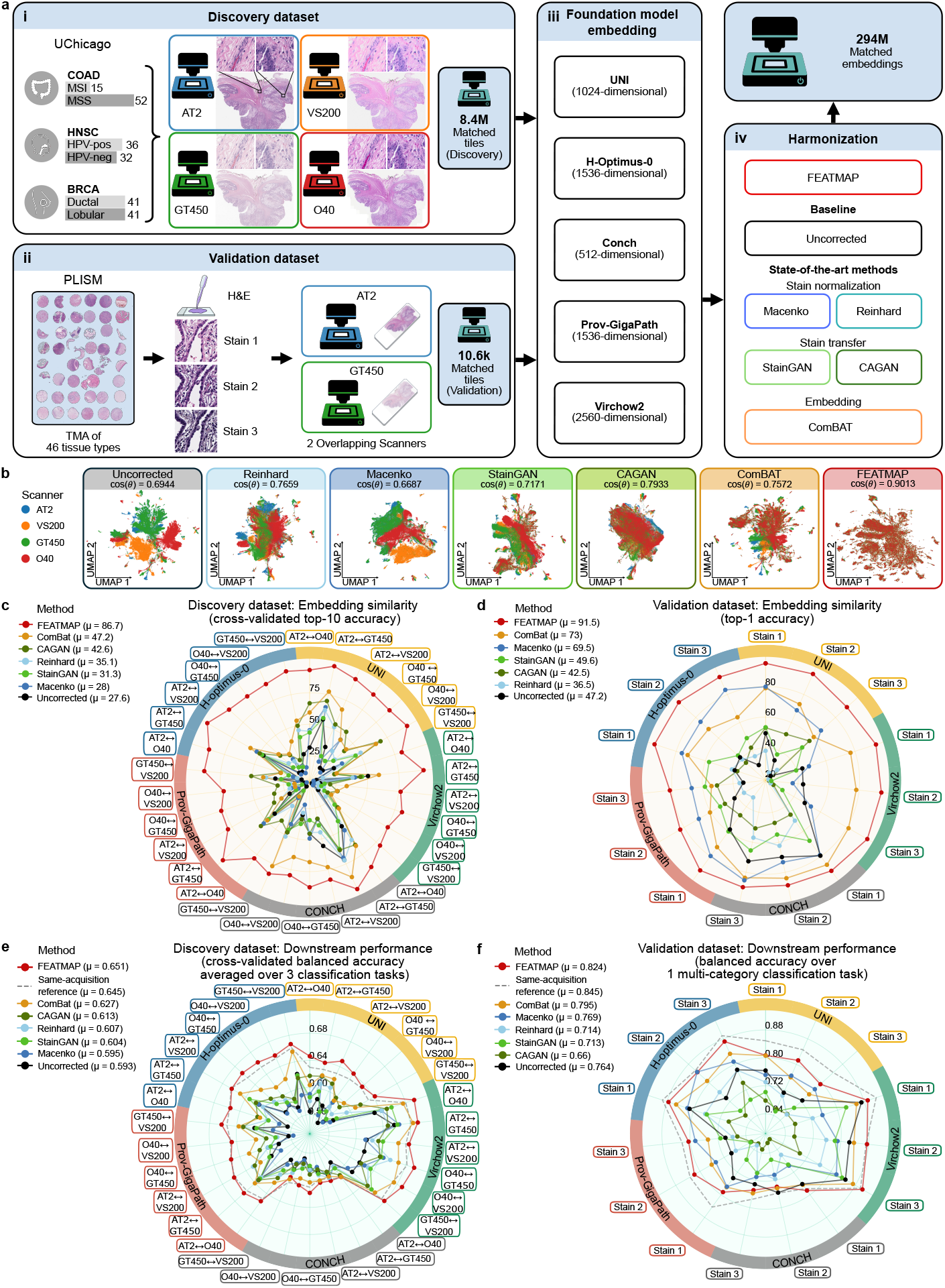
| FEATMAP harmonizes across scanner hardware in digital pathology. **a**, Scanner harmonization study design. The UChicago discovery dataset contains ∼ 8.4M matched tile embeddings from slides scanned on four scanners (i); the PLISM validation dataset contains ∼ 10.6k matched tiles from two overlapping scanners across 46 tissue types and three staining protocols (ii). Both datasets were encoded with five foundation models (iii). FEATMAP was compared with an uncorrected baseline, Reinhard [12], Macenko [13], StainGAN [32], CAGAN [33], and ComBat [34] (iv). **b**, UMAPs of scanner-specific embedding distributions in the discovery dataset, colored by scanner, with average cross-scanner cosine similarity above each plot. **c**,**d**, Embedding similarity in discovery and validation, measured by cross-validated Top-10 accuracy, across scanner pairs and foundation models in discovery and Top-1 accuracy stratified by staining protocol and foundation model in validation. **e**,**f**, Clinical downstream performance in discovery and validation, measured by cross-validated balanced accuracy across three tasks and all scanner pairs for discovery and stratified by staining protocol and foundation model in validation. Dashed lines indicate the same-scanner training/testing reference, which serves as an idealized reference point for the expected upper performance limit. Legend values report method averages (*µ*) across scanners, staining protocols, and foundation models.

We evaluated embedding similarity using Top-k accuracy, defined as the percentage of cases in which a tile’s matched counterpart (same biological content, different scanner) appears among the k nearest neighbors within the full embedding set. We confirmed our findings over additional similarity metrics yielding consistent performance patterns, including cosine similarity (Supplementary Fig. 1) and additional Top-k accuracy values with different k (Supplementary Fig. 2). Since the absolute values of Top-k accuracy depend on the total number of comparisons, which differs between the discovery and validation cohorts, we selected different values of k to suit each dataset’s size.

Consistent with the embedding similarity, UMAP visualizations of the internal embedding distributions reveal severe domain stratification driven by scanner rather than biology (Fig. 2b). Color-based normalization methods (Reinhard, Macenko, StainGAN, CAGAN) reduced scanner separation but produced visible manifold collapse, yielding more homogenized clusters with potential loss of biological substructure. In contrast, embedding-level methods (including FEATMAP) integrated scanner domains while preserving the topological complexity of the embedding space, confirming the ability to harmonize domain shifts without sacrificing biological signal. We additionally proved the ability of FEATMAP to suppress acquisition signatures through a scanner recoverability test in the discovery and validation cohorts (Supplementary Fig. 4).

In the discovery dataset, FEATMAP consistently produced the most similar corrected embeddings across every pairwise combination of scanner and foundation model (Fig. 2c). In contrast, the other correction methods showed little improvement over the uncorrected baseline. Among these methods, CAGAN and ComBat achieved the highest performance, but their gains were small and inconsistent across scanner–foundation model pairs. While performance variation between foundation models was minimal, distinct scanner-specific signatures were apparent. Notably, embeddings from the AT2 and GT450 scanners showed high baseline pairwise similarity despite visible differences in tissue coloration. Since both scanners are manufactured by Leica, this suggests that shared hardware features, rather than image coloration, play a larger role in influencing the embedding structure. External validation on the PLISM dataset mirrored the same trends as in the internal dataset: FEATMAP remained the most robust method for cross-scanner alignment regardless of staining protocol (Fig. 2d) and tissue type (Supplementary Fig. 3). CAGAN, the stain transfer method with the best reported image-level color correction metrics [33], suffered substantial performance degradation on the external cohort, falling below the uncorrected baseline.

Although embedding similarity indicates robust harmonization, clinical downstream model performance represents the most important metric for evaluating foundation model generalization across different scanners. We conducted cross-scanner generalization experiments using four clinical prediction tasks that have been previously accomplished using deep learning for digital pathology: microsatellite stability prediction in colorectal cancer [42], HPV status prediction in head and neck squamous cell carcinoma [43], histological subtyping in breast cancer [44] (UChicago discovery cohort), and a tissue classification task [45] (PLISM validation cohort). For each method, attention-based multiple instance learning (ABMIL) classification models were trained on embeddings from a single source scanner and evaluated on hold-out cases from different scanners using repeated cross-validation, mimicking real-world deployment in which test data originates from different scanners.

Downstream performance was quantified using balanced accuracy to account for class imbalance (and AUC in Supplementary Fig. 5), with same-scanner training and validation serving as same-acquisition reference ceiling (dashed line in Fig. 2e). In the discovery evaluation, most harmonization strategies yielded performance comparable to or only marginally better than the uncorrected baseline, whereas FEATMAP consistently maximized predictive accuracy across all tasks (Fig. 2e, red line). Notably, FEATMAP frequently surpassed the same-acquisition reference, which serves as the theoretical performance ceiling. External validation on the PLISM dataset confirmed these findings (Fig. 2f). Whereas embedding similarity was dominated by scanner-specific effects with marginal differences between foundation models, downstream performance revealed clear stratification between foundation models following previously reported performance rankings [46], but negligible scanner-pair differences (Supplementary Fig. 6 shows balanced accuracy and AUC in all cohorts).

### 2.2 FEATMAP harmonizes across the digital pathology foundation model domain

We investigated whether the same paired-data framework could align embedding spaces generated by different digital pathology foundation models. In this experiment, the foundation model type was treated as the primary acquisition domain rather than scanner type. We used the same discovery and validation datasets as in the scanner harmonization experiment, but extracted embeddings from each tile using five different foundation models (Fig. 2a). Each pathology tile therefore yielded the paired five foundation model embeddings with identical biological content, allowing FEATMAP to isolate foundation model-specific signatures.

Because existing harmonization methods are not designed to map between foundation model embedding spaces with different dimensionalities, we evaluated FEATMAP against an uncorrected baseline. This design allowed us to test whether FEATMAP could learn robust cross-foundation-model mappings despite differences in embedding size, architecture, and training history.

In the embedding similarity the Top-10 accuracy of the uncorrected baseline nears zero across all pairwise combinations in both discovery and validation evaluations (Fig. 3b), and UMAP visualization confirms complete separation of foundation model-specific clusters with little observable overlap (cos(*θ*) = 0.0000; Fig. 3a). This is due to the significant challenge of harmonizing different foundation models. Whereas different scanners introduce relatively subtle shifts within a single embedding space, different foundation models produce embeddings in entirely disjoint representation spaces that may differ in dimensionality, training objective and learned feature hierarchy. Despite this extreme starting point, FEATMAP substantially recovered cross-foundation-model embedding similarity (Fig. 3a). In the discovery dataset, FEATMAP achieved a mean Top-10 accuracy of 73.5 across all 20 pairwise foundation model combinations (Fig. 3b). Performance varied across foundation model pairs, with some combinations proving easier to align than others, likely reflecting varying degrees of shared embedding structure between foundation models as well as correlating to similar embedding dimensionality. External validation on the PLISM dataset confirmed generalization of the learned mappings (Fig. 3c), though absolute Top-10 values were lower than in the discovery dataset. We confirmed that our findings were robust across additional similarity metrics, with consistent performance patterns observed for cosine similarity (Supplementary Fig. 7), additional Top-k accuracy values (Supplementary Fig. 8), and across different tissue types (Supplementary Fig. 9). The UMAP visualization after FEATMAP harmonization shows the effect of the transformation, wherein foundation model-specific clusters merge into a single integrated manifold with a cosine similarity of 0.9361 (Fig. 3a), demonstrating that FEATMAP recovers a shared geometric structure across embedding spaces while preserving biological variation.

**Fig. 3.**
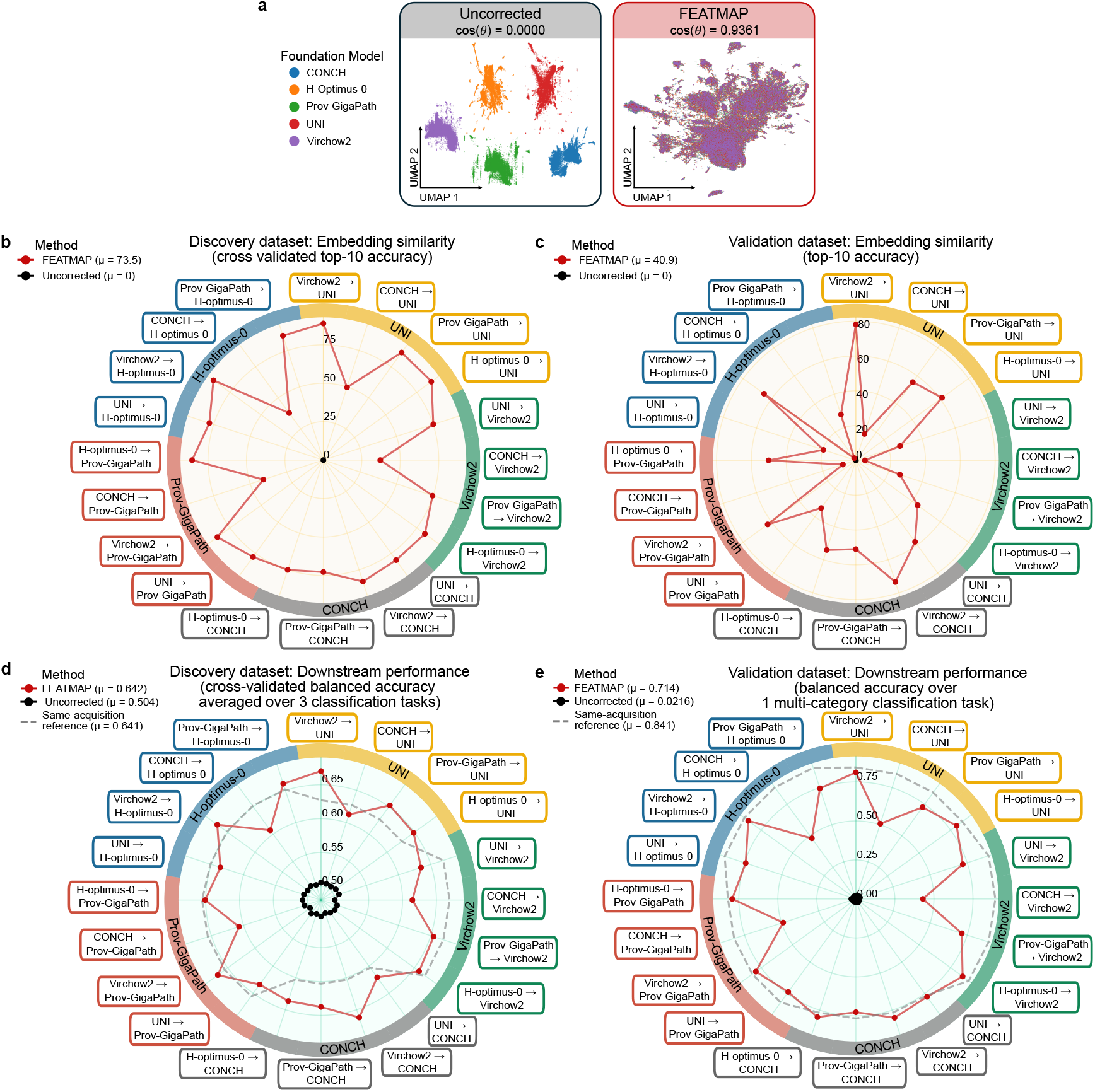
| FEATMAP harmonizes across foundation models in digital pathology. **a**, UMAP visualizations of embedding distributions colored by foundation model before and after harmonization, with average cosine similarity indicated above each plot. Uncorrected embeddings form distinct model-specific clusters, whereas FEATMAP harmonization into the smallest embedding space (CONCH) integrates embeddings from different foundation models into a shared manifold while preserving biological substructure. **b**, Quantitative evaluation of cross-model embedding alignment in the discovery dataset using cross-validated Top-10 accuracy across all 20 ordered pairwise foundation-model combinations among UNI, H-Optimus-0, CONCH, Prov-GigaPath, and Virchow2. FEATMAP is compared with the uncorrected baseline only, because no existing harmonization method directly addresses cross-foundation-model alignment. **c**, External validation of cross-model embedding alignment using Top-10 accuracy in the validation dataset. **d**, Downstream clinical prediction performance in the discovery cohort was evaluated as cross-validated balanced accuracy under a cross-foundation-model generalization protocol in which downstream models are trained on embeddings from one foundation model and tested on embeddings from a different foundation model. **e**, Downstream clinical prediction performance in the validation dataset was evaluated using balanced accuracy. Dashed lines indicate the same-foundation-model training/testing reference, which serves as an idealized reference point for the expected upper performance limit. Legend values report method averages (*µ*) across scanners and staining protocols.

To evaluate whether this improved embedding alignment translates to better results in a clinical downstream task, we assessed downstream prediction performance using the same cross-condition generalization protocol used for scanner harmonization measured in balanced accuracy (Fig. 3d, 3e) and AUROC (Supplementary Fig. 10) . ABMIL classification downstream models were trained on embeddings from one foundation model and evaluated on embeddings from a different foundation model, with same-acquisition performance (same-model training and testing) serving as the idealized reference point for the expected upper performance limit. The uncorrected baseline yielded zero accuracy across all foundation model pairs (Fig. 3d), confirming that foundation model embeddings cannot be compared without harmonization. FEATMAP closed this gap (*µ* = 0.714), recovering most of same-acquisition performance (*µ* = 0.841) across all pairwise combinations, scanners (Supplementary Fig. 11) and clinical cohorts (Supplementary Fig. 12). External validation mirrored these trends (Fig. 3e), with FEATMAP consistently approaching the same-acquisition reference.

### 2.3 FEATMAP harmonizes across the brain MRI field strength domain

To assess whether FEATMAP generalizes beyond digital pathology, we applied it to brain MRI, where scanners operating at different magnetic field strengths (1.5T and 3T) introduce systematic intensity and contrast differences that propagate into foundation model embeddings. We trained FEATMAP on 918 matched T1-weighted scan pairs from the human brain phantom dataset [47], in which the same individual was scanned repeatedly on 1.5T and 3T machines (Fig. 4a i). Scans underwent standardized preprocessing (Fig. 4a iii) before embedding with BrainIAC [48], a brain MRI foundation model producing a 768-dimensional embedding space (Fig. 4a iv). The learned transformation was then evaluated on the Yale longitudinal dataset of brain metastases on MRI [49], comprising 3,906 matched T1-scan pairs from patients scanned at both field strengths (Fig. 4a ii). FEATMAP was benchmarked against an uncorrected baseline, four image-level normalization methods (histogram equalization – HE, adaptive histogram equalization – AHE, contrast-limited adaptive histogram equalization – CLAHE, and Z-score normalization), as well as the embedding-level method ComBat (Fig. 4a v).

**Fig. 4.**
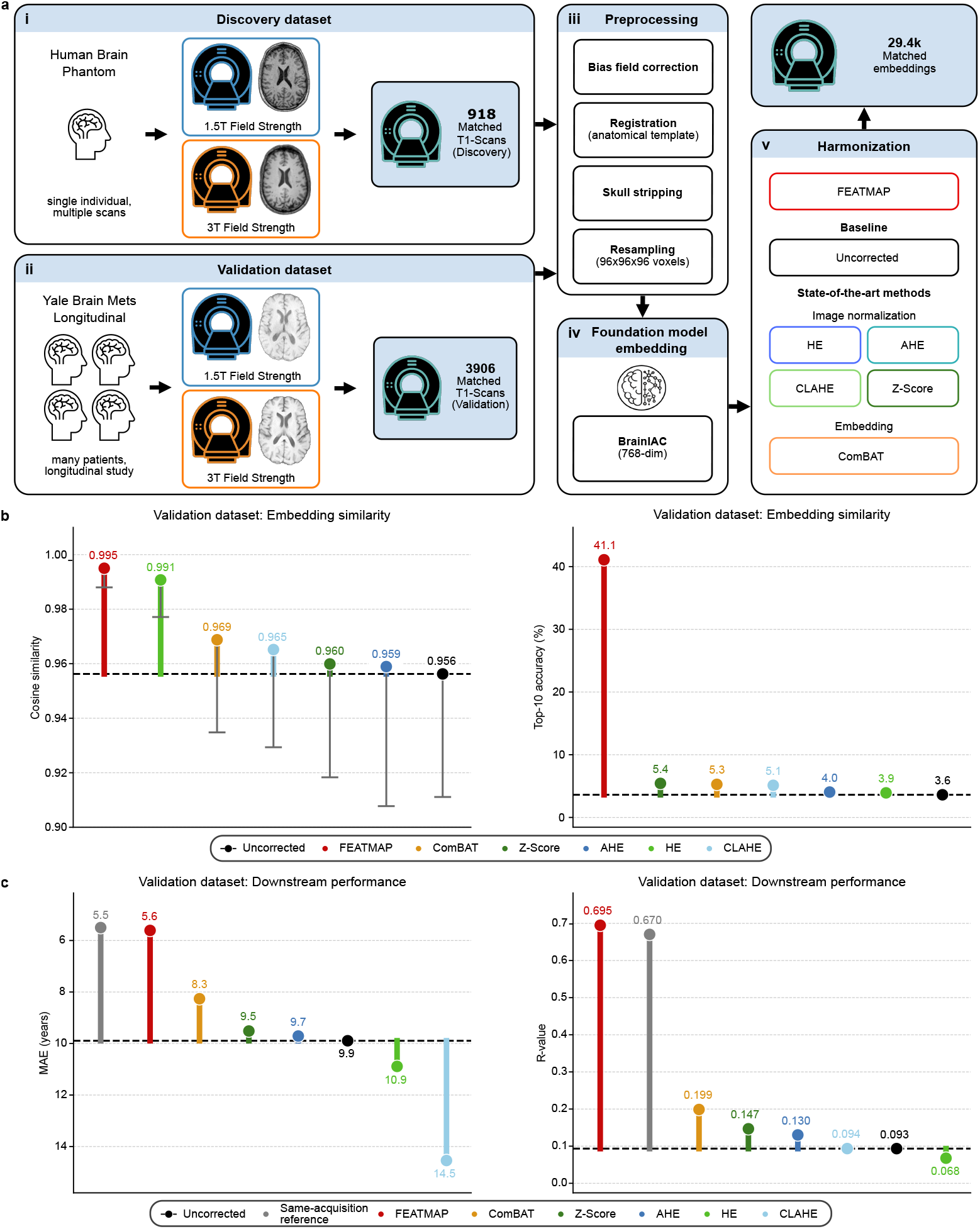
| FEATMAP harmonizes brain MRI embeddings across field strengths. **a**, MRI field strength harmonization study design. The discovery dataset includes 918 matched T1-weighted brain phantom scan pairs acquired at 1.5T and 3T (i), and the external Yale Brain Mets Longitudinal validation dataset includes 3,906 matched T1-weighted scan pairs from a multi-patient cohort (ii). Scans were preprocessed (iii), then embedded with BrainIAC to obtain 768-dimensional representations (iv). FEATMAP was compared with an uncorrected baseline, four image-level normalization methods—histogram equalization (HE), adaptive histogram equalization (AHE), contrast-limited AHE (CLAHE), and Z-score normalization—and embedding-level ComBat (v). **b**, External validation of cross-field-strength embedding alignment using cosine similarity between matched scan pairs and Top-10 accuracy. The cosine-similarity y-axis is cropped to highlight method differences. **c**, External downstream performance on brain-age prediction under a cross-field-strength setting, with models trained on one field strength and evaluated on the other, with performance measured using MAE (lower is better) and R-value (higher is better). The MAE y-axis is inverted and cropped to highlight differences. Same-field-strength training/testing is shown as an idealized upper-performance reference.

Validation results revealed superior performance of FEATMAP compared to other methods. In embedding similarity, cosine similarity values were uniformly high across all methods compared to the previous harmonization experiments, reflecting the relatively subtle nature of field strength shifts compared to the dramatic domain gaps in scanner or foundation model harmonization (Fig. 4b, left). However, Top-10 accuracy exposed larger differences because, as a relative retrieval metric, it amplifies small shifts in embedding similarity that absolute similarity metrics can smooth over. FEATMAP achieved a Top-10 accuracy of 41.1, whereas all other methods remained near baseline levels (range 2.9–5.4; Fig. 4b, right). This dissociation illustrates why distance-based metrics alone can be misleading for evaluating harmonization, since small systematic embedding shifts can prevent correct nearest-neighbor retrieval despite high overall cosine similarity. Only FEATMAP corrected these shifts effectively. This distinction is clinically relevant because downstream models may rely on subtle embedding differences as predictive signals, which may explain the observed performance gap.

To evaluate downstream clinical utility, we assessed cross-field-strength generalization on a brain age prediction task using only the validation dataset, since the discovery dataset only covered a single individual within two years. Downstream models were trained on embeddings from one field strength and evaluated on the other, with downstream performance measured using mean absolute error (MAE) and Pearson correlation (R-value).

The uncorrected baseline yielded an MAE of 9.9 and an R-value of 0.093, reflecting substantial degradation relative to the same-acquisition reference (MAE = 5.5, R = 0.670). Image-level normalization methods (HE, AHE, CLAHE, Z-score) performed comparably to or worse than the uncorrected baseline (MAE range 9.5–14.5, R range 0.068–0.147). CLAHE suffered severe degradation (MAE = 14.5, R = 0.094), indicating that excessive contrast manipulation is counterproductive. ComBat offered a slight improvement over the baseline (MAE = 8.3, R = 0.199). In contrast, FEATMAP recovered to almost the same-acquisition performance (MAE = 5.6, R = 0.695) across both metrics, reducing the cross-condition performance gap to a minimum.

These results demonstrate that FEATMAP’s harmonization strategy transfers effectively from digital pathology to radiology, despite fundamental differences in data modality, foundation model architecture, and the nature of the acquisition signature.

## 3 Discussion

Medical foundation models compress complex biomedical inputs into high-dimensional embeddings that support diverse downstream clinical tasks. However, reliable cross-institutional comparability of these embeddings and the models derived from them remains challenging, because embeddings can encode acquisition signatures that are unrelated to underlying biology. FEATMAP provides a scalable, modular, and targeted framework for correcting specific acquisition signatures in medical foundation model embeddings. Across three distinct harmonization settings, FEATMAP substantially increased cross-condition embedding similarity and reduced downstream performance gaps, demonstrating that targeted geometric correction can improve the interoperability of foundation model representations.

Existing approaches to harmonization have typically operated either at the input level, by applying normalization before embedding, or at the embedding level, by adjusting features without explicitly distinguishing acquisition signatures from biological variation. Both strategies have shown limited efficacy. FEATMAP addresses these limitations by learning a targeted, reusable affine transformation directly in the embedding space from paired data in which biological content is held constant while the acquisition condition is systematically varied. This design aims to preserve the biological and demographic variation that constitutes the signal of interest for downstream clinical tasks. Because each transformation is learned independently for a given domain shift, multiple transformations can be applied sequentially, for example first correcting for scanner model and then for staining protocol. This composability follows from the affine structure and distinguishes FEATMAP from approaches that require joint re-estimation when new sources of variation are introduced. In the MRI analysis, field strength may be confounded with scanner model, site, acquisition protocol, reconstruction pipeline, and scan time. Therefore, the observed harmonization effect should be interpreted as correction of field-strength-associated acquisition variation rather than the isolated effect of field strength.

The ability of a single global affine transformation to harmonize embeddings between acquisition signatures suggests that these signatures are stable patterns seperable from other underlying representational structure. Rather than fundamentally reorganizing the embedding space, acquisition signatures appear to induce smooth geometric distortions between similarly arranged low-dimensional subspaces. Singular value decomposition of the learned transformations further showed that their dominant component was rotational, supporting the interpretation that FEATMAP primarily aligns embedding spaces through structured geometric reorientation. The field-strength harmonization experiments reinforce these interpretations: FEATMAP’s strategy transferred effectively from digital pathology to radiology despite fundamental differences in data modality and in the nature of the acquisition signature. Moreover, the paired phantom data, consisting of repeated acquisitions of the same medical specimen under different conditions, proved sufficient to calibrate harmonization for an entire clinical cohort. This finding is relevant for institutions seeking to pool retrospective imaging data acquired with heterogeneous acquisition parameters.

These findings also raise broader questions about the local geometry of foundation model embedding spaces. While existing theoretical frameworks suggest that foundation model spaces may be organized globally as collections of lower-dimensional manifolds within high-dimensional representation spaces, the internal geometric shape of these manifolds remains poorly understood. One possible interpretation is that, if semantically related observations within a manifold are approximately distributed according to high-dimensional Gaussian-like structure, they may concentrate near thin shell-like regions, as predicted by the thin shell phenomenon [50]. This view is consistent with empirical observations of radial or “onion-like” organization in embedding spaces [19]. It may also explain why Gaussian or second-order alignment can be beneficial [51], yet insufficient to fully remove acquisition-specific distortions, because real embedding manifolds are unlikely to be perfectly Gaussian and may instead form distorted shell-like structures with residual non-Gaussian geometry.

The foundation model harmonization results carry implications beyond practical interoperability. The observation that a learned affine mapping recovers substantial cross-model embedding compatibility is consistent with the Platonic Representation Hypothesis [28]. Specifically, medical foundation models of the same type (such as pathology foundation models) trained on overlapping data, but with different architectures and objectives, converge toward shared representational geometries that are largely recoverable through structured alignment. At the same time, the residual gap between FEATMAP-harmonized and same-acquisition performance indicates that models retain architecture-specific information not captured by a single global affine map. This points toward model-specific refinements or higher-order corrections as directions for further improvement. FEATMAP may also serve as a computationally efficient initialization for constructing joint embedding spaces across models, reducing representational differences through constrained linear alignment before fine-tuning with methods such as contrastive learning that can capture residual nonlinear structure.

An unexpected finding across multiple settings was that FEATMAP-harmonized embeddings occasionally exceeded the same-acquisition reference in downstream performance. One possible explanation is that the affine constraint acts as a form of beneficial regularization, projecting embeddings onto a lower-dimensional or less noisy manifold and thereby facilitating more robust convergence in downstream models. Understanding this regularization effect more precisely, and determining when it reliably holds, may help improve downstream frameworks such as ABMIL.

FEATMAP has two main limitations. First, it requires paired calibration data in which biological content is held constant while acquisition conditions vary, which may be difficult to obtain for certain acquisition conditions. Second, it models the whole harmonization with a single global affine transformation, which may underfit local, subpopulation-specific effects. This limitation is consistent with the Aristotelian Representation Hypothesis [52], which emphasizes shared local neighborhood structure rather than convergence toward a single global representation. Thus, FEATMAP is most feasible when standardized calibration material can be repeatedly measured, and future extensions could use local, class-conditional, or region-specific transformations, although these would require more data and add modeling complexity.

The current formulation also targets one domain shift at a time and assumes that different acquisition effects can be treated as approximately independent. Sequential composition provides a practical way to combine transformations, but interactions between domains, such as scanner-specific staining effects or protocol-dependent field-strength artifacts, may require joint modeling. Finally, although FEATMAP generalized across the settings evaluated here and provides a theoretical basis for expecting similar behavior in analogous cases, its effectiveness for acquisition signatures with fundamentally different characteristics remains to be tested empirically. Future work could extend the framework to account for class-conditional or region-specific shifts, to other modalities such as spatial omics or multimodal imaging, and to residual nonlinear domain effects.

FEATMAP’s closed-form estimation and modular design make it well suited for integration into clinical deployment pipelines as a lightweight preprocessing step. In summary, FEATMAP demonstrates that principled geometric assumptions about foundation model embedding spaces can yield harmonization tools that are simple, interpretable, and effective.

## 4 Methods

### 4.1 UChicago dataset

To study scanner-induced domain variation, we collected whole-slide images (WSIs) from three cohorts digitized on four distinct brightfield pathology scanners: Leica Aperio AT2, Olympus Slideview VS200, Leica Aperio GT 450 and Grundium Ocus 40. All slides represented identical tissue sections captured under different scanner devices. The three cohorts are head and neck squamous cell carcinoma (HNSC) with 68 patients, invasive breast carcinoma (BRCA) with 82 patients and colon adenocarcinoma (COAD) with 67 patients, each with one representative WSI per patient. Clinical labels used for downstream analyses were HPV status in HNSC (36 HPV-positive, 32 HPV-negative), tumor subtype in BRCA (41 ductal, 41 lobular) and microsatellite status in COAD (15 MSI, 52 MSS). For tile alignment, slides were initially aligned manually to the closest global match at three reference locations, after which tiles were refined automatically using phase-correlation-based registration and verified manually by inspecting matched tiles at 50 random locations per WSI. This procedure yielded approximately 8.4 million pixel-perfect aligned tile pairs across all four scanners. This dataset served as the discovery cohort for learning the FEATMAP scanner and foundation-model transformations.

### 4.2 PLISM dataset

For external validation in digital pathology, we used a subset of the PLISM Tile Dataset [40, 41], comprising pre-aligned tiles from consecutive tissue microarray (TMA) slides stained across 13 H&E staining conditions and scanned on seven pathology scanners. To reduce complexity, we selected the three stains most similar to routine H&E for evaluation [53]: GV – Stain 1, GVH – Stain 2, and GMH – Stain 3. The dataset includes 46 tissues from diverse anatomical sites, with two scanners overlapping those in our discovery dataset, enabling cross-institutional validation across tissue types. Tissue type was used as the label for the downstream classification task. This yielded approximately 10.6k matched tile pairs across the two overlapping scanners. This dataset served as the validation cohort for the scanner and foundation-model harmonization settings.

### 4.3 Human brain phantom dataset

To study field-strength-induced domain variation in brain MRI, we used the human brain phantom dataset [47] in which a single individual was scanned repeatedly on 1.5T and 3T MRI machines. We utilized scans from comparable machines acquired at 1.5T and 3T that shared the same bore diameter, manufacturer, and system architecture platform, and were acquired within 1 year of each other. All eligible scans at each field strength were paired combinatorially, so that each 1.5T scan was matched with every qualifying 3T scan, allowing for multiple pairs. This yielded 918 matched T1-weighted scan pairs. Because all scans originate from the same healthy individual within a short time period, biological variation is held constant and any systematic differences in the resulting embeddings can be attributed to the field-strength acquisition condition. This dataset served as the discovery cohort for the field-strength harmonization setting.

### 4.4 Yale Brain Mets Longitudinal dataset

For the external validation of the findings in brain MRI using the human “brain phantom” dataset, we used a subset of the Yale Brain Mets Longitudinal dataset [49], a longitudinal study in which we selected individuals who were unintentionally scanned on both field strengths over time. As with the discovery dataset, we selected scans from comparable machines at 1.5T and 3T that shared the same bore diameter, manufacturer, and system architecture platform, and were acquired within one year of each other to reduce temporal biological variation. All eligible scans at each field strength were paired combinatorially, so that each 1.5T scan was matched with every qualifying 3T scan, permitting multiple pairs per patient, which results in non-independent scan pairs. This yielded 3,906 matched T1-weighted scan pairs. Because the dataset includes many patients with diverse pathologies, it enables evaluation of whether the transformation learned on a single-subject phantom dataset generalizes to a heterogeneous clinical population. This dataset served as the validation cohort for the field-strength harmonization setting.

### 4.5 Digital pathology preprocessing and embedding extraction

Aligned whole-slide images were tiled into non-overlapping 128 *µ*m ×128 *µ*m patches (256 × 256 pixels at 20 × magnification) using Slideflow [54]. Each tile was embedded using five pathology foundation models: UNI [35] (1,024 dimensions), H-Optimus-0 [36] (1,536 dimensions), Conch [37] (512 dimensions), Prov-GigaPath [38] (1,536 dimensions), and Virchow2 [39] (2,560 dimensions). All models were applied with their published default preprocessing and inference settings. The same tiling and embedding procedure was applied to both the UChicago and PLISM datasets.

### 4.6 Brain MRI preprocessing and embedding extraction

All T1-weighted MRI scans underwent a standardized preprocessing pipeline. First, bias field correction was applied. Scans were then registered to a standard anatomical template [48] to establish spatial correspondence across subjects and field strengths. Skull stripping was performed to remove non-brain tissue, and the resulting brain volumes were resampled to a uniform resolution of 96 × 96 × 96 voxels. Preprocessed scans were embedded using BrainIAC [48], a brain MRI foundation model producing 768-dimensional representations. The same preprocessing and embedding pipeline was applied to both the human brain phantom discovery dataset and the Yale Brain Mets Longitudinal validation dataset (Fig. 4).

### 4.7 FEATMAP

FEATMAP operates on the observation that foundation model embeddings are subject to non-biological acquisition conditions (such as scanner hardware, model architecture, or MRI field strength) that introduce systematic shifts unrelated to the underlying biology that we call acquisition signatures. We denote the set of acquisition domains by *D* = {scanner, architecture, field strength, … } . For each acquisition domain *d* ∈ *D*, we define a finite set of conditions *C*_*d*_ that specifies the choices within that acquisition domain; for example, *C*_Scanner_ = { AT2, GT450, VS200, O40} . Not all acquisition conditions need to be known for a given sample, and a given sample is generally subject to a composition of conditions across multiple acquisition domains.

Each biological sample is represented by a collection of instance-level embeddings extracted from the raw input data using a foundation model. These embeddings correspond to localized elements of the sample, such as image tiles in digital pathology or volumetric patches in MRI. For a sample with *T* instance-level embeddings of dimension *k*, we stack the embeddings row-wise to form a matrix *Z* ∈ ℝ^*T ×k*^ . Because our paired-data design holds biology constant while varying the acquisition conditions within a single acquisition domain *d*, the resulting embedding matrices lie in a acquisition-domain-specific feature space 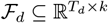, where *T*_*d*_ reflects the number of instance-level embeddings associated with acquisition domain *d*.

For a given acquisition domain *d* with source condition *c* and target condition *c*^*′*^, FEATMAP learns a transformation 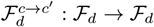 that maps embeddings from condition *c* into condition *c*^*′*^. Each such transformation is parameterized as a learned affine map:

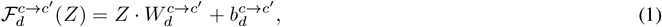

where 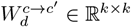 is the linear component and 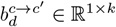 is the translation. The linear component can be interpreted as a combination of elementary geometric operations in the embedding space, including rotation, scaling, and shear, although FEATMAP does not explicitly enforce such a decomposition. When transforming between differently sized feature spaces, *W* is rectangular rather than square. The parameters *W* and *b* are estimated from paired observations across conditions (for example identical tissue sections scanned on two different scanners) using ordinary least-squares regression that minimizes the squared error between the transformed source embeddings and the target embeddings. This admits a closed-form solution, making FEATMAP a transparent estimation procedure that requires no iterative or black-box optimization.

Because all transformations within one acquisition domain act on the common feature space *F*_*d*_, they can be composed. For three conditions *c, c*^*′*^, *c*^*′′*^ ∈ *C*_*d*_, the mapping from *c* to *c*^*′′*^ via the intermediate condition *c*^*′*^ is given by

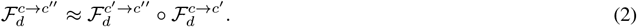

Since each component is affine, the composition is also affine:

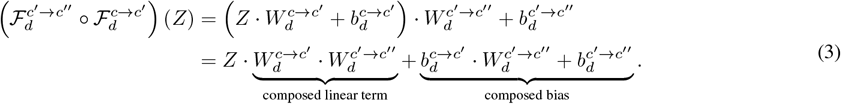

This composability extends beyond acquisition conditions and into acquisition domains. Given a sequence of acquisition domains *d*_1_, …, *d*_*m*_ ∈ *D*, alignment is performed by successive acquisition domain transformations

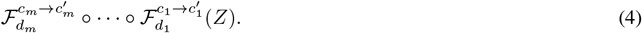

For example, one can first harmonize across model architectures and then across scanners by applying

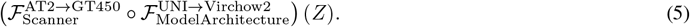

This modular structure allows new acquisition domains to be added without re-estimating existing transformations, and each transformation can be stored and reused across tasks and cohorts.

### 4.8 Scanner harmonization experiment

For scanner harmonization, the acquisition domain was defined by scanner model (*s* = Scanner), with four conditions corresponding to the four pathology scanners in the UChicago dataset. We compared FEATMAP against six widely used harmonization strategies, including two image-based color normalization methods, Reinhard [12] and Macenko [13], two image-based stain transfer models, StainGAN [32] and CAGAN [33], and one embedding-level harmonization method, ComBat [34]. All methods were evaluated alongside an uncorrected baseline. Image-based methods were applied to all WSIs before embedding extraction with the respective foundation model, whereas embedding-level methods, ComBat and FEATMAP, were applied after embedding generation. In the UChicago discovery cohort, FEATMAP was evaluated using a two-fold cross-validation strategy in which pairwise affine transformations were estimated using one half of the patients in the discovery data and then applied to the held-out half. For external validation in the PLISM cohort, FEATMAP was trained on the entire UChicago discovery dataset and subsequently applied to the PLISM validation dataset without retraining. ComBat was applied directly using the original implementation [34] to the full UChicago discovery dataset and the PLISM validation dataset independently.

### 4.9 Foundation model harmonization experiment

To test whether FEATMAP generalizes to model-induced variation, we treated differences among the five foundation models as the acquisition domain (*d* = Model Architecture) rather than scanner model. In the UChicago discovery cohort, we used the same two-fold cross-validation strategy as for scanner harmonization, in which FEATMAP was trained to estimate pairwise affine transformations using embeddings from one half of the patients and then applied to the held-out half. For external validation in the PLISM cohort, FEATMAP was trained on the entire UChicago discovery dataset and subsequently applied to the PLISM validation dataset without retraining. Because no existing method is designed to harmonize between foundation model embedding spaces of different dimensionality, FEATMAP was benchmarked solely against an uncorrected baseline, in which the embedding spaces were made compatible by appropriate vector padding.

### 4.10 Field-strength harmonization experiment

For brain MRI harmonization, the acquisition domain was defined by magnetic field strength (*d* = Field Strength), with two conditions corresponding to 1.5T and 3T MRI acquisitions. We compared FEATMAP against five commonly used harmonization strategies, including four image-level normalization methods, histogram equalization (HE), adaptive histogram equalization (AHE), contrast-limited adaptive histogram equalization (CLAHE), and Z-score normalization, and one embedding-level harmonization method, ComBat. All methods were evaluated alongside an uncorrected baseline. Image-level methods were applied to the preprocessed MRI volumes before embedding extraction, whereas embedding-level methods, ComBat and FEATMAP, were applied after embedding generation. FEATMAP was trained on the 918 matched scan pairs from the human brain phantom discovery dataset and subsequently validated on the 3,906 matched pairs from the Yale Brain Mets Longitudinal dataset without retraining. ComBat was applied directly using the original implementation [34] to the human brain phantom discovery dataset and the Yale Brain Mets Longitudinal validation dataset independently.

### 4.11 Embedding similarity metrics

We evaluated harmonization quality using acquisition-signature-aligned paired data. For each paired instance *i*, we computed cosine similarity between the harmonized source embedding and its paired target embedding:

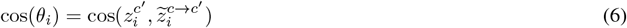

where 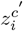 is the embedding under the target condition *c’* and 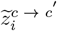 is the harmonized source embedding. We summarized cosine similarity by taking the mean across all instances. We additionally computed Top-k accuracy (k = 1, 5, 10), defined as the percentage of cases in which a matched counterpart (same biological content, different acquisition condition) appears among the k nearest neighbors within the full embedding set. Top-k accuracy was chosen as the primary metric because raw distance measures, such as cosine similarity, can misleadingly reward methods that reduce biologically meaningful variation by compressing all embeddings toward a homogeneous distribution.

### 4.12 Downstream prediction performance metrics

For digital pathology, we trained attention-based multiple instance learning (ABMIL) classification models on harmonized embeddings derived from a single source condition and evaluated performance on hold-out cases from different conditions using 50 times repeated stratified 5-fold cross-validation. Performance was quantified using balanced accuracy and area under the receiver operating characteristic curve (AUROC) to account for class imbalance. Same-condition training and validation served as the same-acquisition reference (performance ceiling). In the UChicago cohort, the three clinical prediction tasks were microsatellite stability prediction in colorectal cancer, HPV status prediction in HNSC and histological subtyping in BRCA. In the PLISM cohort, the downstream task was tissue type classification across 46 anatomical sites.

For brain MRI, downstream performance was evaluated on a brain age prediction task. Models were trained on embeddings from one field strength and evaluated on the other. Performance was quantified using mean absolute error (MAE) and Pearson correlation coefficient (R-value), with the same-acquisition performance serving as the reference.

### 4.13 Scanner recoverability

We evaluated attention-based multiple instance learning (ABMIL) classification models on the raw and harmonized embeddings to predict the scanner model using 50 times repeated stratified 5-fold cross-validation and quantified performance using balanced accuracy. Results are averaged across foundation models and for the discovery dataset across the three cohorts.

## Supplementary Information

**Supplementary Fig. 1.**
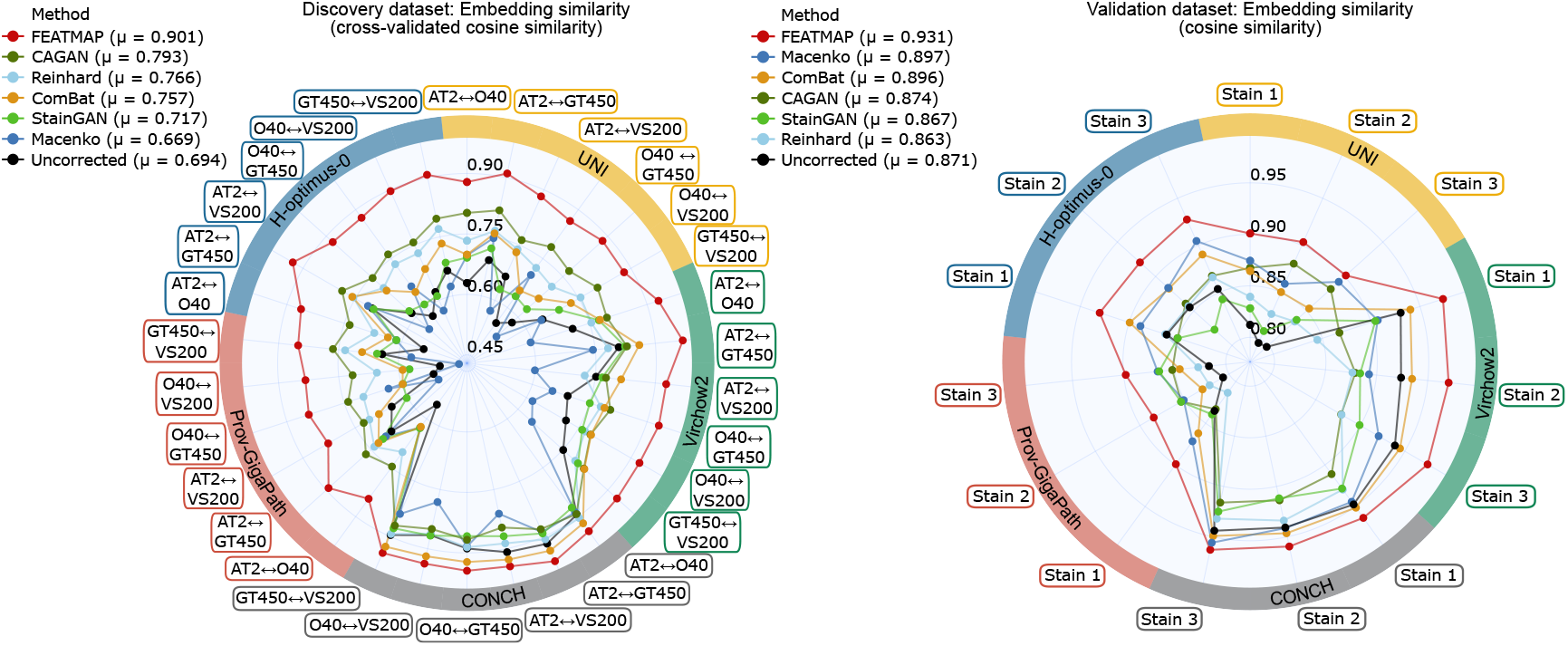
Cosine similarity analysis supporting the scanner harmonization experiments in the discovery dataset (left) and validation dataset (right). Legend values report method averages (*µ*) across scanners, staining protocols, and foundation models.

**Supplementary Fig. 2.**
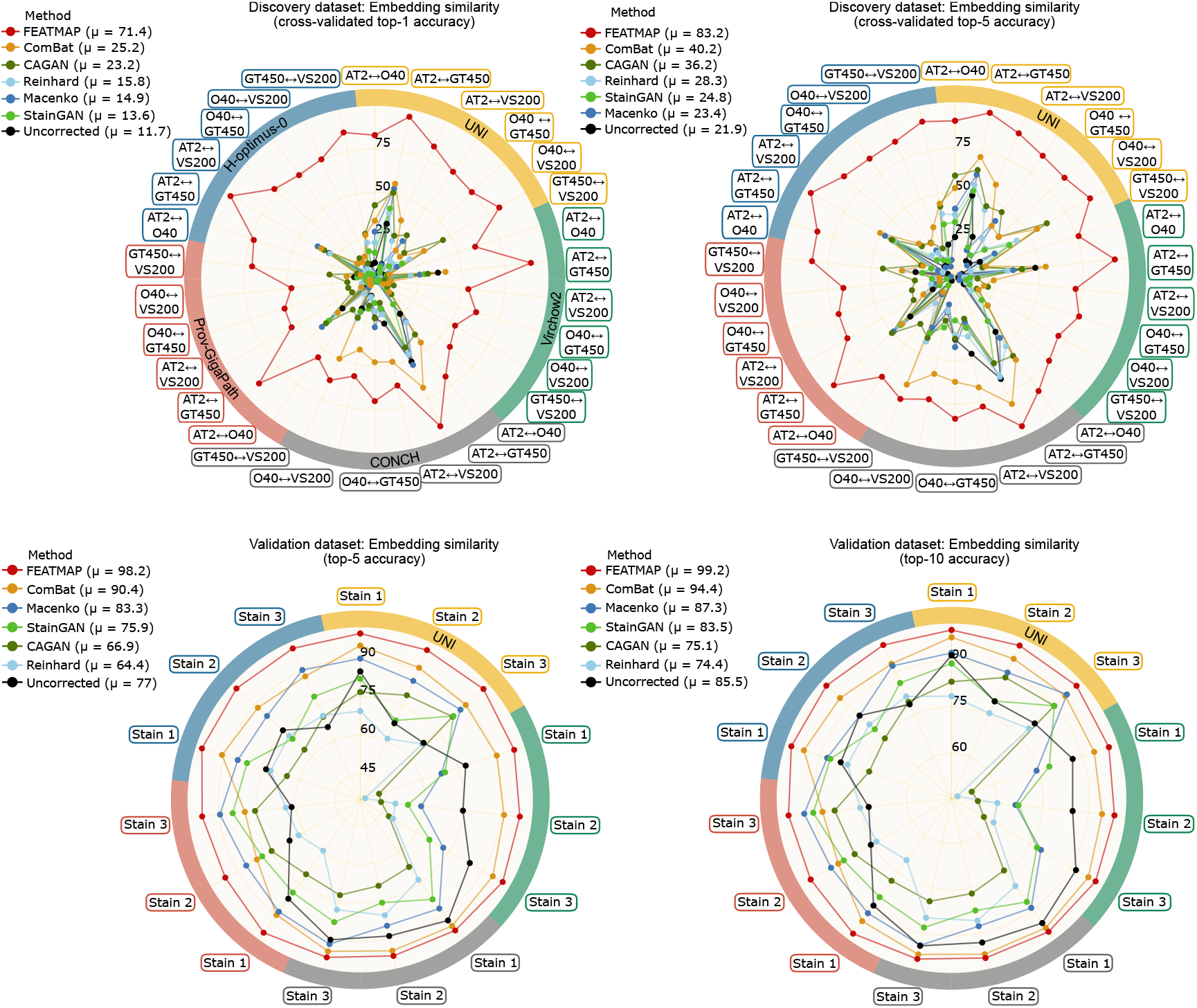
Additional Top-k accuracy results supporting the scanner harmonization experiments. Cross-validated Top-1 accuracy (top left) and Top-5 accuracy (top right) are shown for the discovery dataset, and Top-5 accuracy (bottom left) and Top-10 accuracy (bottom right) are shown for the validation dataset. Legend values report method averages (*µ*) across scanners, staining protocols, and foundation models.

**Supplementary Fig. 3.**
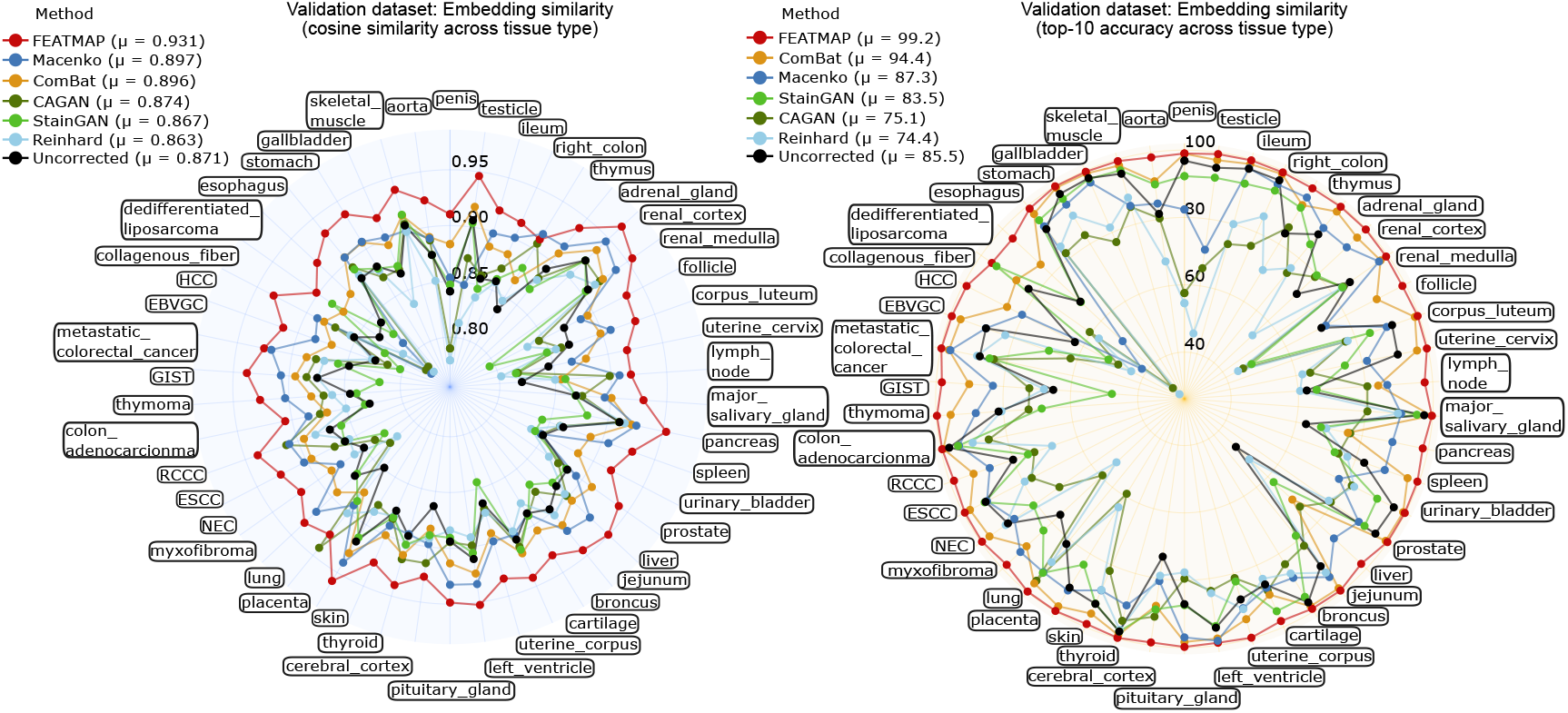
Tissue-type-stratified embedding similarity results supporting the scanner harmonization experiments in the validation dataset, measured using cosine similarity (left) and Top-10 retrieval accuracy (right). Legend values report method averages (*µ*) across tissue types.

**Supplementary Fig. 4.**
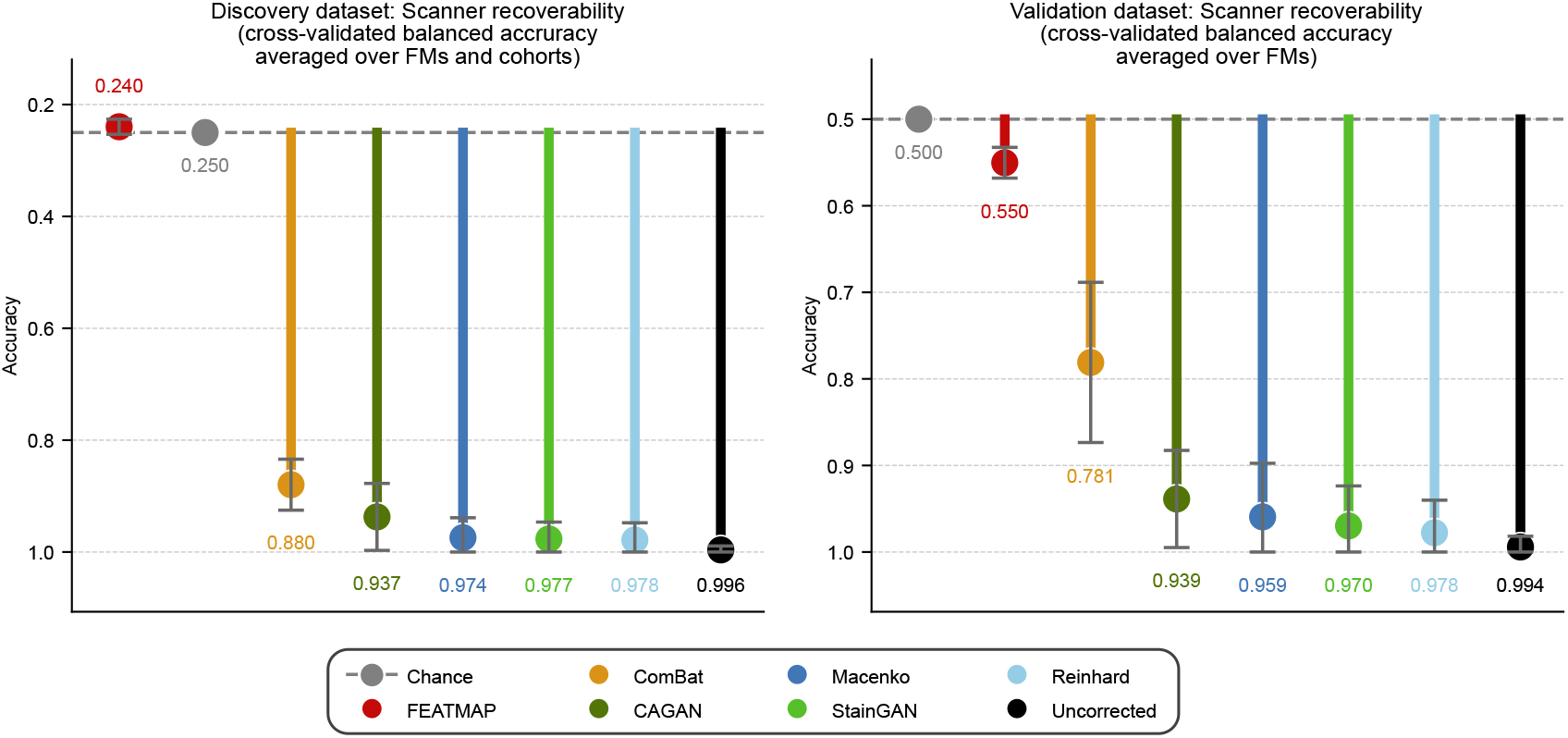
Scanner recoverability analysis supporting the scanner harmonization experiments. Scanner classification performance was evaluated using cross-validated balanced accuracy in the discovery dataset (left) and validation dataset (right). Dashed lines indicate chance-level performance with 25% for the four-class scanner classification task in the discovery dataset and 50% for the binary scanner classification task in the validation dataset, which both serve as a theoretical reference point for the expected performance limit.

**Supplementary Fig. 5.**
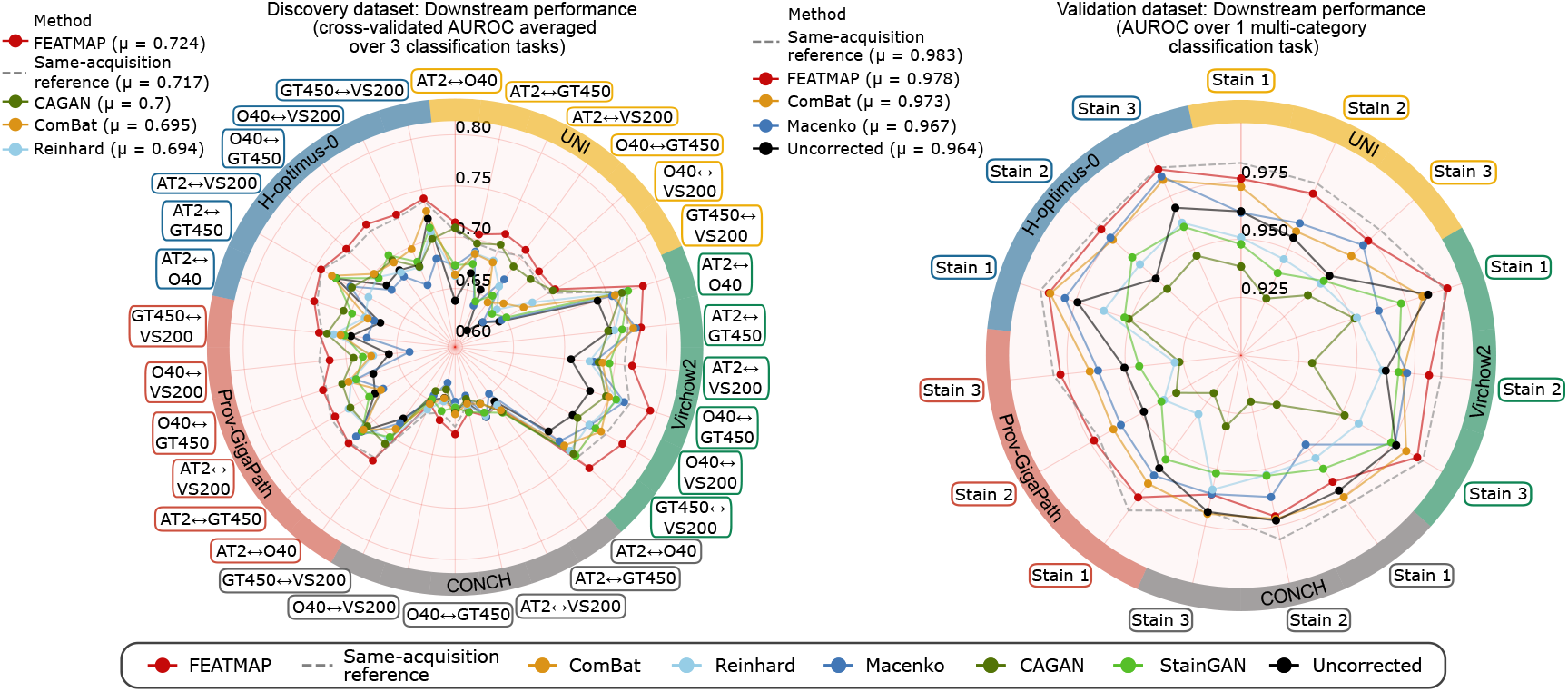
Downstream performance analysis supporting the scanner harmonization experiments, measured using AUROC in the discovery dataset (left) and validation dataset (right). Dashed lines indicate the same-scanner training/testing reference, which serves as an idealized reference point for the expected upper performance limit. Legend values report method averages (*µ*) across scanners, staining protocols, and foundation models.

**Supplementary Fig. 6.**
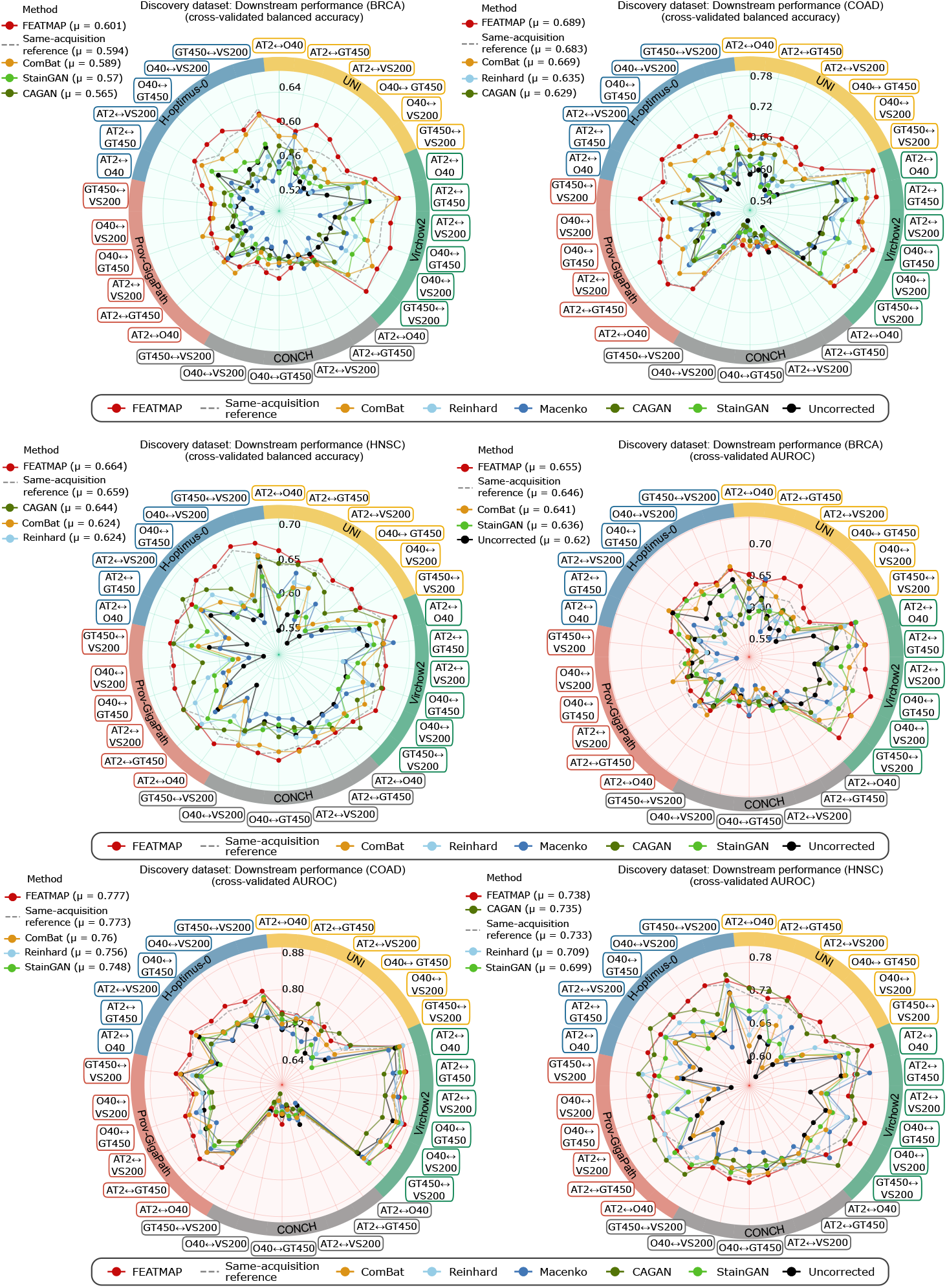
Cohort-specific downstream performance supporting the scanner harmonization experiments in the discovery dataset. Balanced accuracy is shown for the BRCA (top left), COAD (top right), and HNSC (middle left) cohorts, and AUROC is shown for the BRCA (middle right), COAD (bottom left), and HNSC (bottom right) cohorts. Dashed lines indicate the same-scanner training/testing reference, which serves as an idealized reference point for the expected upper performance limit. Legend values report method averages (*µ*) across scanners, staining protocols, and foundation models.

**Supplementary Fig. 7.**
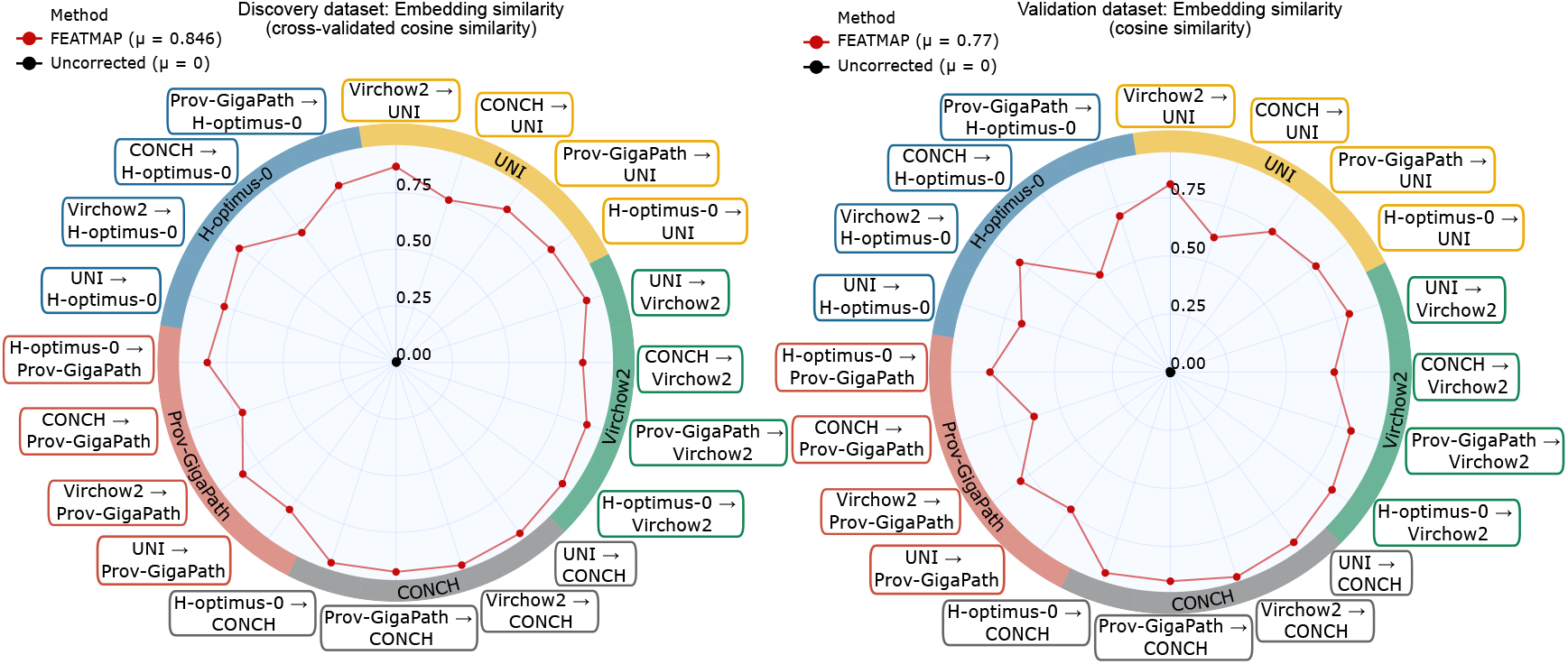
Cosine similarity analysis supporting the foundation-model harmonization experiments in the discovery dataset (left) and validation dataset (right). Legend values report method averages (*µ*) across foundation models.

**Supplementary Fig. 8.**
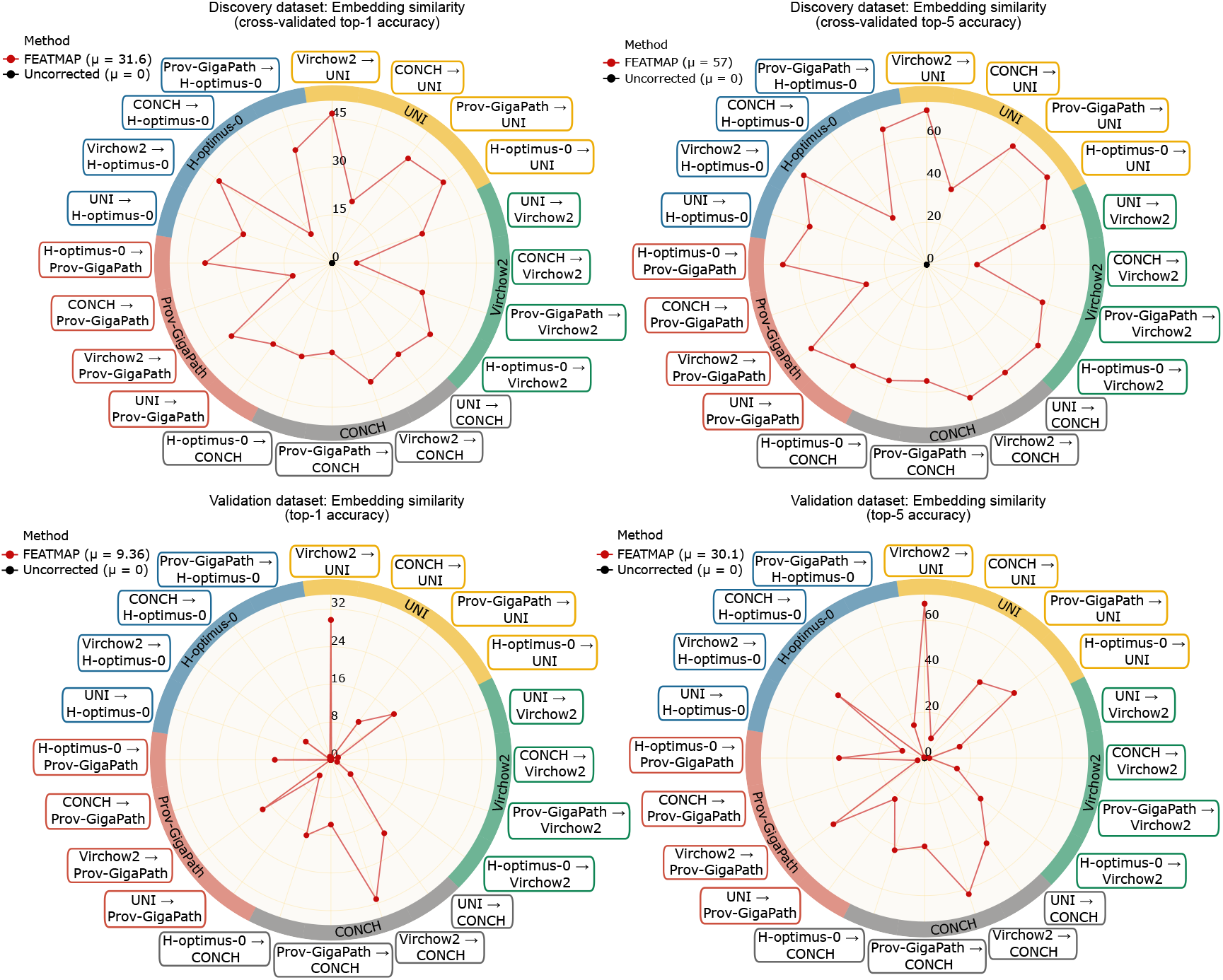
Additional Top-k accuracy results supporting the foundation-model harmonization experiments. Cross-validated Top-1 accuracy (top left) and Top-5 accuracy (top right) are shown for the discovery dataset, and Top-1 accuracy (bottom left) and Top-5 accuracy (bottom right) are shown for the validation dataset. Legend values report method averages (*µ*) across foundation models.

**Supplementary Fig. 9.**
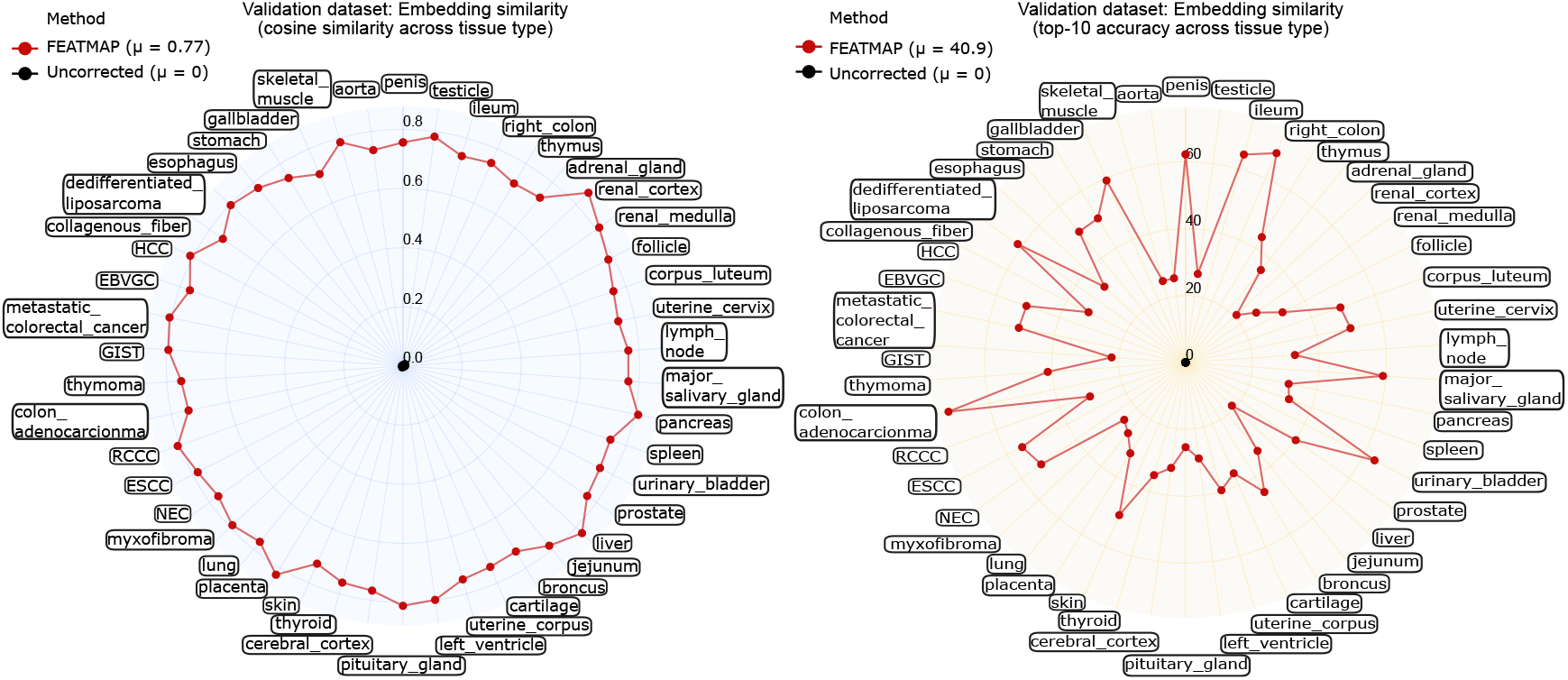
Tissue-type-stratified embedding similarity results supporting the foundation-model harmonization experiments in the validation dataset, measured using cosine similarity (left) and Top-10 retrieval accuracy (right). Legend values report method averages (*µ*) across foundation models.

**Supplementary Fig. 10.**
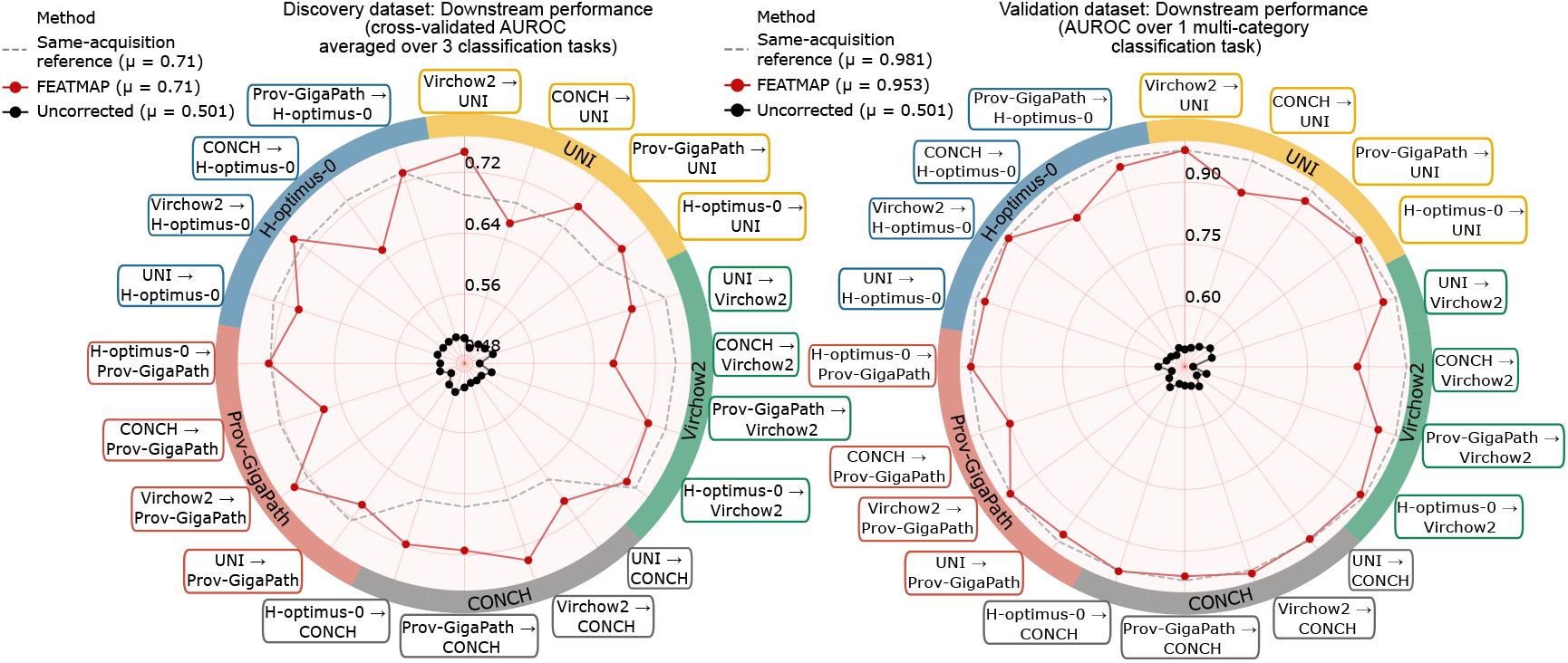
Downstream performance analysis supporting the foundation-model harmonization experiments, measured using AUROC in the discovery dataset (left) and validation dataset (right). Dashed lines indicate the same-foundation-model training/testing reference, which serves as an idealized reference point for the expected upper performance limit. Legend values report method averages (*µ*) across foundation models.

**Supplementary Fig. 11.**
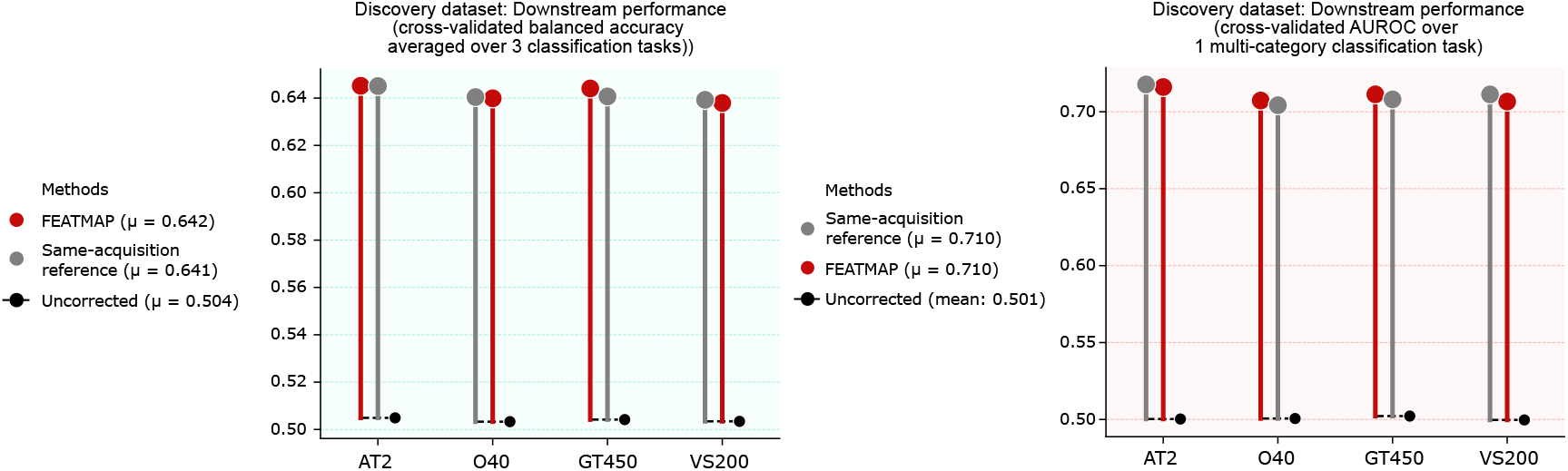
Scanner-stratified downstream performance supporting the scanner harmonization experiments in the discovery dataset. Results are shown across the four scanners and averaged over foundation models, measured using balanced accuracy (left) and AUROC (right). Grey points indicate the same-foundation-model training/testing reference, which serves as an idealized reference point for the expected upper performance limit. Legend values report method averages (*µ*) across scanners.

**Supplementary Fig. 12.**
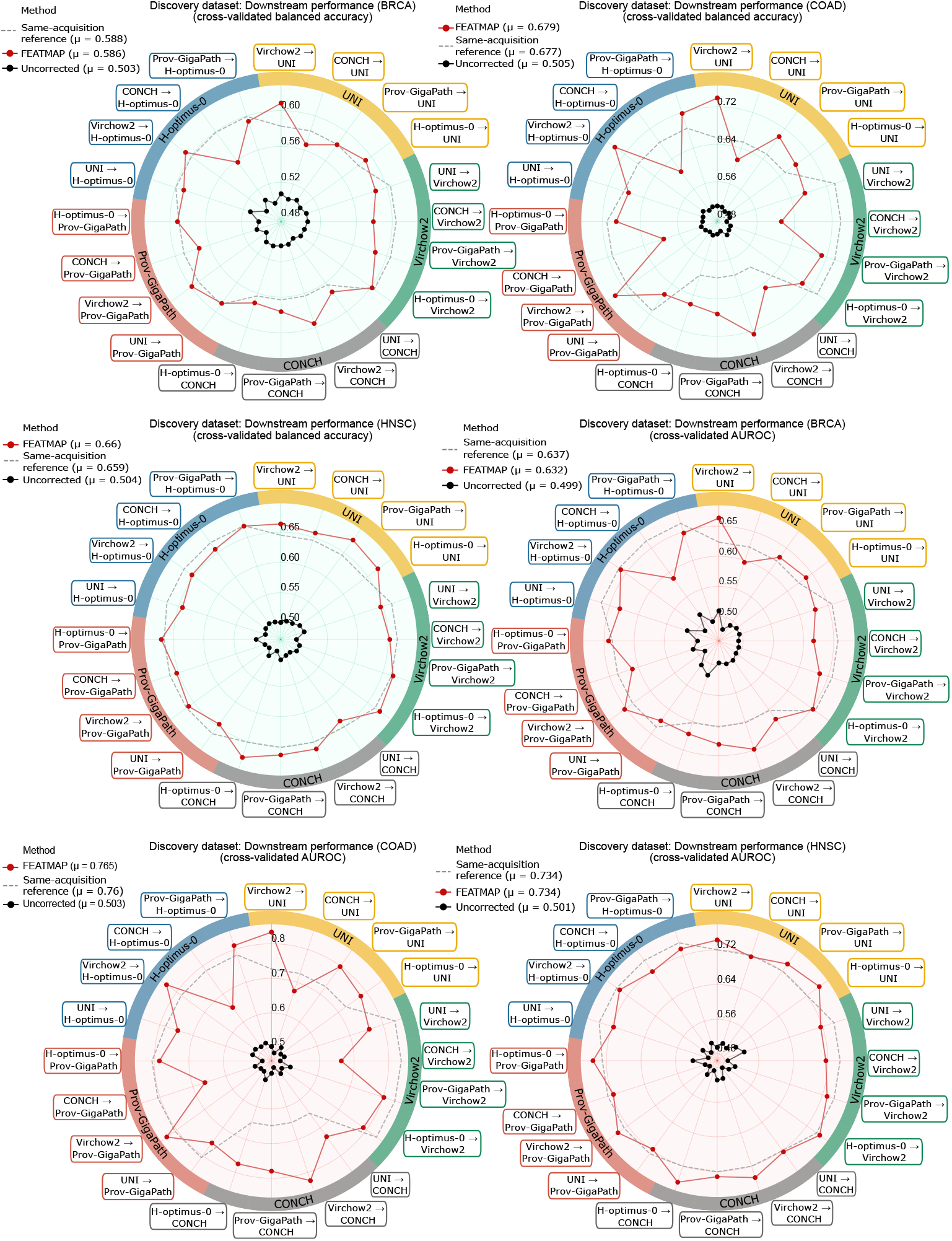
Cohort-specific downstream performance supporting the foundation-model harmonization experiments in the discovery dataset. Balanced accuracy is shown for the BRCA (top left), COAD (top right), and HNSC (middle left) cohorts, and AUROC is shown for the BRCA (middle right), COAD (bottom left), and HNSC (bottom right) cohorts. Dashed lines indicate the same-foundation-model training/testing reference, which serves as an idealized reference point for the expected upper performance limit. Legend values report method averages (*µ*) across foundation models.

